# Precentral gyrus abnormal connectivity in male and female schizophrenia patients

**DOI:** 10.1101/143768

**Authors:** Mahdi Zarei

**Author notes:** Electronic address.

## Abstract

The present study demonstrates the Precentral gyrus (PreCG) alteration and its abnormal connectivities in Schizophrenia (SCH) patients. The resting-sate functional magnetic resonance imaging (rs-fMRI) data of healthy control subjects and SCH patients (Centre for Biomedical Research Excellence data set) are used to examine the aberrant functional brain connectome in SCH which contains raw anatomical and functional MR data from 72 patients with Schizophrenia and 75 healthy controls, ranging in age from 18 to 65 years old. Our results show that PreCG has abnormal communication with Thalamus, Hippocampus, Parahippocampal Gyrus (pPaHC), posterior division of Supramarginal Gyrus (pSMG) and medial prefrontal cortex (mPFC) (p-FDR=0.05).

## 1 Introduction

Recent studies shows that Precentral gyrus (PreCG) has significant reduced of functional activity (FC) in Schizophrenia (SCH) patients [16, 13]. Also, SCH is associated with volume deficits in Precentral gyrus [15]. Another research indicates that the patients with SCH showed lower activation in left PreCG than right PreCG [10] and a regional homogeneity (ReHo) study showed decreased ReHo in right Precentral gyrus [19].

In this research we examined the PrecG functional connectivity impairment in SCH patients using ROI based analysis [14]. We used the Center for Biomedical Research Excellence data set (COBRE) ([8]) to demonstrate how the functional connectivity of the PreCG with the rest of the brain regions changes in SCH. In conjunction with previous studies, our results indicate that the PreCG has abnormal connectivity with some of the brain regions like Thalamus and Hippocampus, but dis impairments are not the similar in two hemisphere. We also analyzed the FC differences between male and female SCH patients and showed that the regions like Thalamus are more affected in the female SCH patients.

## 2 Data

We included 48 subjects from Centre for Biomedical Research Excellence COBRE data set; 24 healthy controls and 24 schizophrenia patients (12 men and 12 women in each group) in the study. In the original data set (see table 2) there are more male subjects than females and to allow the same effects of the both genders in the results we used the above arrangement. COBRE data contains raw anatomical and functional MRI data from patients with Schizophrenia and healthy controls. Some papers that by the COBRE group published based on this data [11, 17, 3]. This data set is available on [8] and contains raw anatomical and functional MR data from patients with Schizophrenia and healthy controls, ranging in age from 18 to 65 years old. Resting fMRI, anatomical MRI, phenotypic data for every participant including: gender, age, handedness and diagnostic information are released.

**Table 1:** Demographic characteristics of subjects in COBRE data [1]

|  | N | Age(SD) | %Female | %Right-Handed |
| --- | --- | --- | --- | --- |
| Schizophrenia | 72 | 38.16(13.89) | 0.19 | 0.83 |
| Patients | 74 | 35.82(11.58) | 0.31 | 0.96 |

**Table 2:** The connectivity contrast values between the PerCG right and the targets (p-value=0.05) (the complete list of the effect size is provided in the supplementary data)

| Analysis Unit | Statistic | p-unc | p-FDR |
| --- | --- | --- | --- |
| Seed Intensity = 18.77, Size = 5 | PreCG r | $F(30)(113) = 3.17$ | 0.0000 |
| PreCG r -Hippocampus l | $T(142) = 4.16$ | 0.0001 | 0.0074 |
| PreCG r -Thalamus l | $T(142) = -3.92$ | 0.0001 | 0.0093 |
| PreCG r -pPaHC l | $T(142) = 3.62$ | 0.0004 | 0.0151 |
| PreCG r -Thalamus r | $T(142) = -3.54$ | 0.0005 | 0.0151 |
| PreCG r -pSMG r | $T(142) = -3.53$ | 0.0006 | 0.0151 |

## 3 Analysis

Different preprocessing methods like realignment, coregistering, normalization are applied for the structural and functional data. The functional volumes are coregistered with the region of interest and structural volumes. Regions of interest and all the Brodmann areas defined from Talairach daemon assigned to all subjects. By segmentation of structural image for each subject, grey matter, white matter and cerebrospinal fluid (CSF) masks generated. Here, the time series of interest are the number of PCA components.

### 3.1 First and second level of covariates

In this step the realignment parameters in BOLD model is defined (first level covariate), then in the second level covariate the group level regressor is performed. We categorized the data input data into 4 groups; Control females, Control males, Patient females and Patient male.

After defining the experiment data, the functional data is imported, then the structural data is segmented to define the grey matter, white matter, cerebrospinal fluid region of interest. By performing PCA on within region of interests the ROIs time series is extracted.

## 4 Denoising of the data

Before analyzing the data we need to explore and remove the confounds. Different source of possible confounds like cerebrospinal fluid and white matter signal and within-subject covariate (realignment parameters) are considered. We chose the 5 dimension that is the number of temporal components are being used. Similarly, the number of dimension for white matter was 5 and the derivative order was 0 and the histogram plot r value before and after confound removal and the band-pass filter is set to [0.008 0.09]. Figure 4 (a and b) shows variance explained (r-square) by each of the possible confounding sources and the histogram plot at the center displays the voxel-to-voxel connectivity (r values) before, and after confound removal for 2 subjects. The histograms r-square in order to identify outlier subject and the quality control is computed for every subject (figure 4 c).

**Figure 1:**
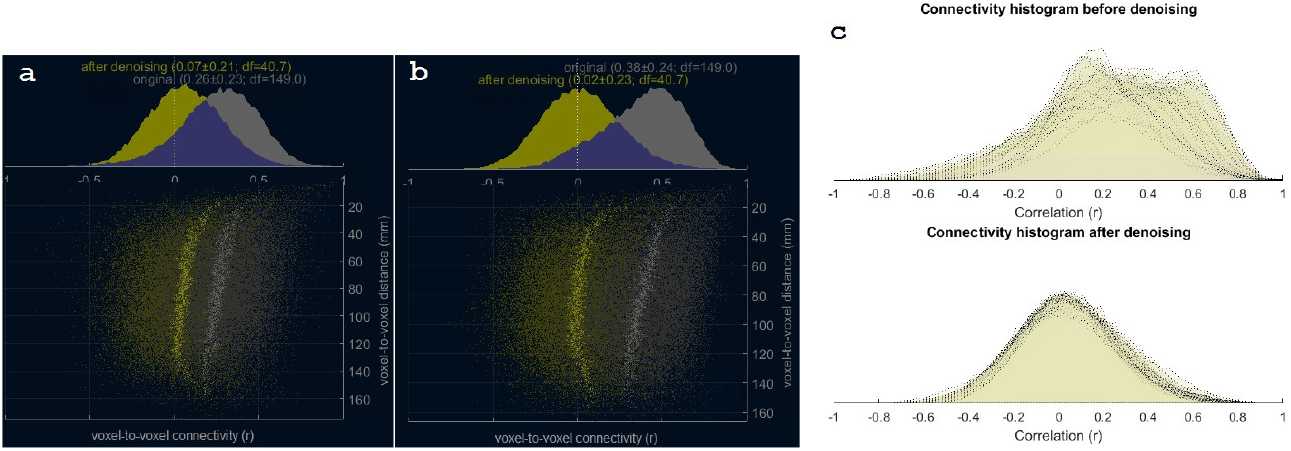
Histogram of functional connectivity values (r). Total variance explained (r-square) by each of the possible confounding sources for 2 subjects.

**Figure 2:**
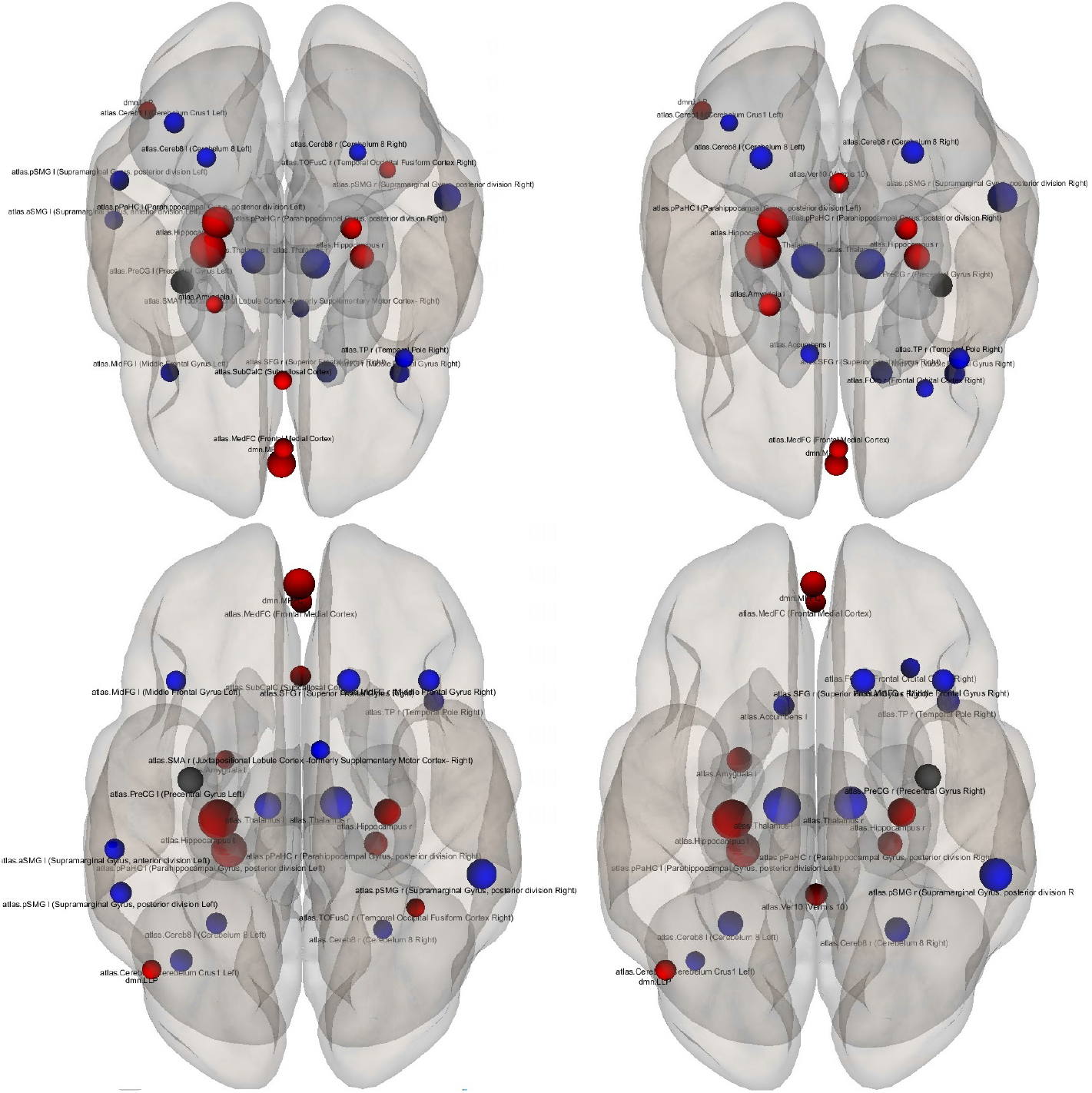
The superior (top) and the posterior views of the ROI-level results for each target ROI. The selected sources (precentral gyrus left and right) are shown by black dots and the targets are blue (control > SCH) and red (control < SCH). Effects size for each region is shown by the dot size.

**Figure 3:**
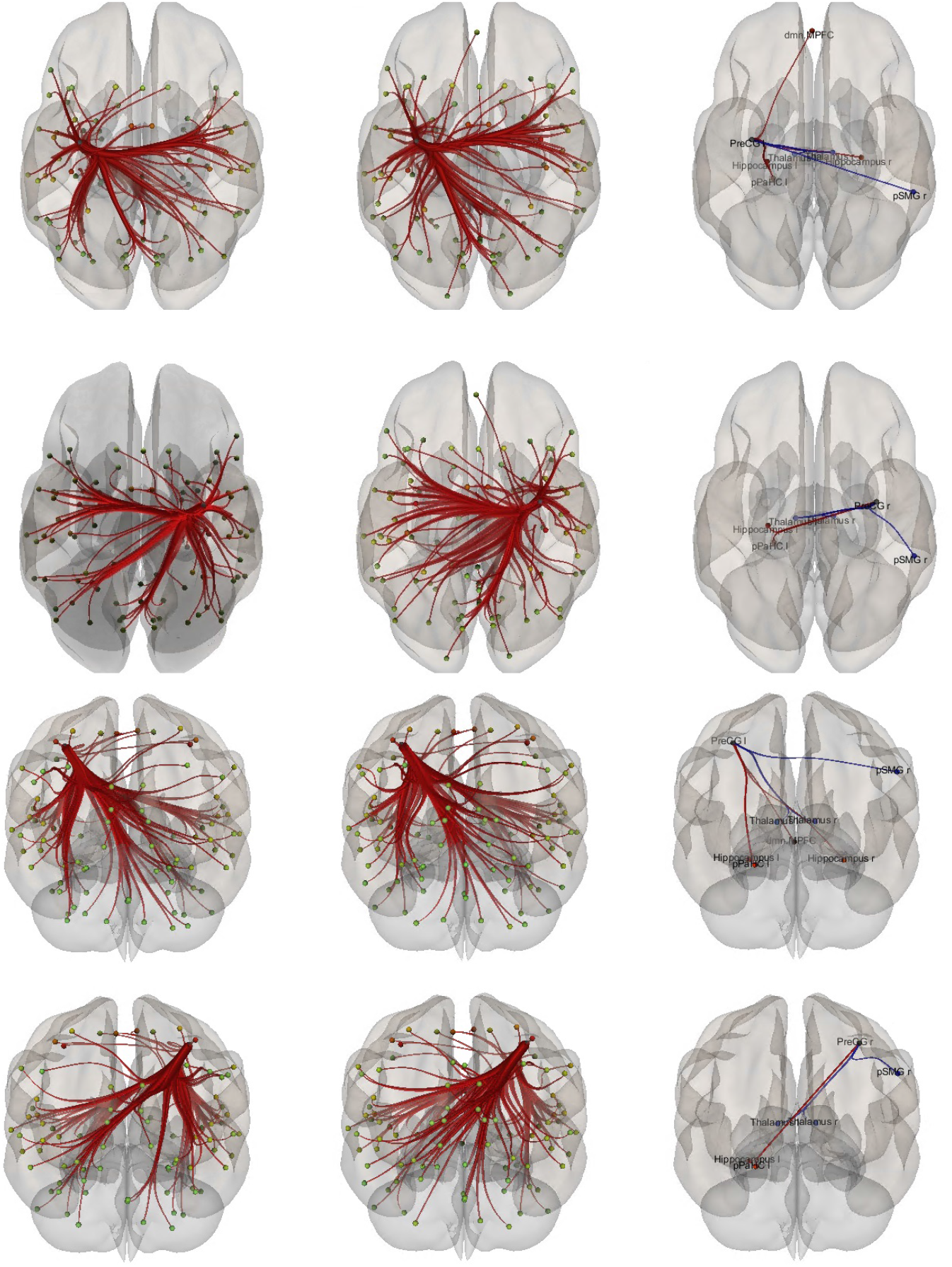
Functional connectivity difference of the left and right precentral gyrus between healthy controls vs. SCH patients (controls>Schizophrenia). The PrecG connectivity inthe healthy control, SCH patients and their difference are shown in the first, second and the last column respectively and the two top raws are the superior views and the the two bottom raws are the posterior views.

**Figure 4:**
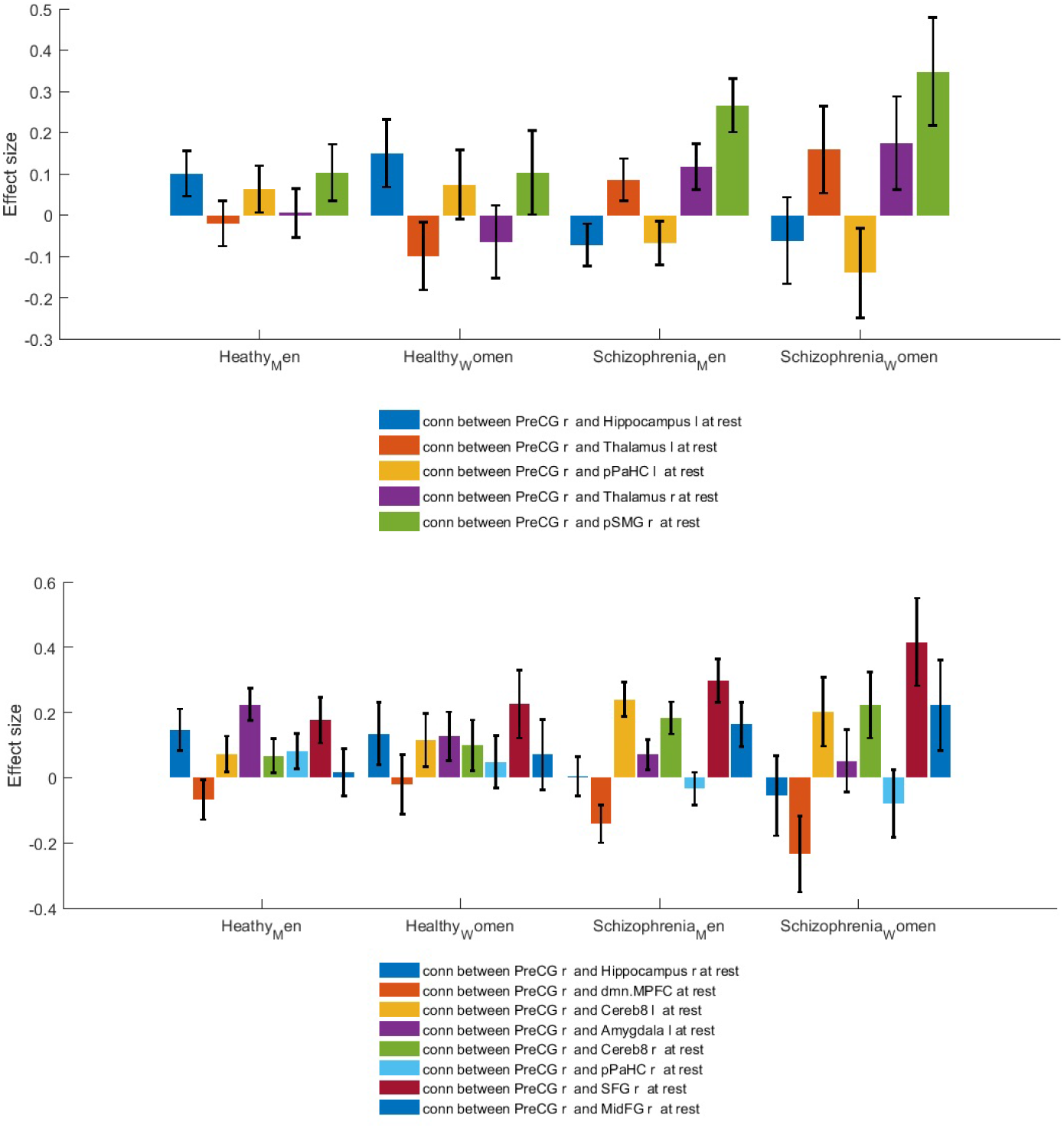
Functional connectivity difference of the … precentral gyrus between healthy controls vs. SCH patients (controls>Schizophrenia).

**Figure 5:**
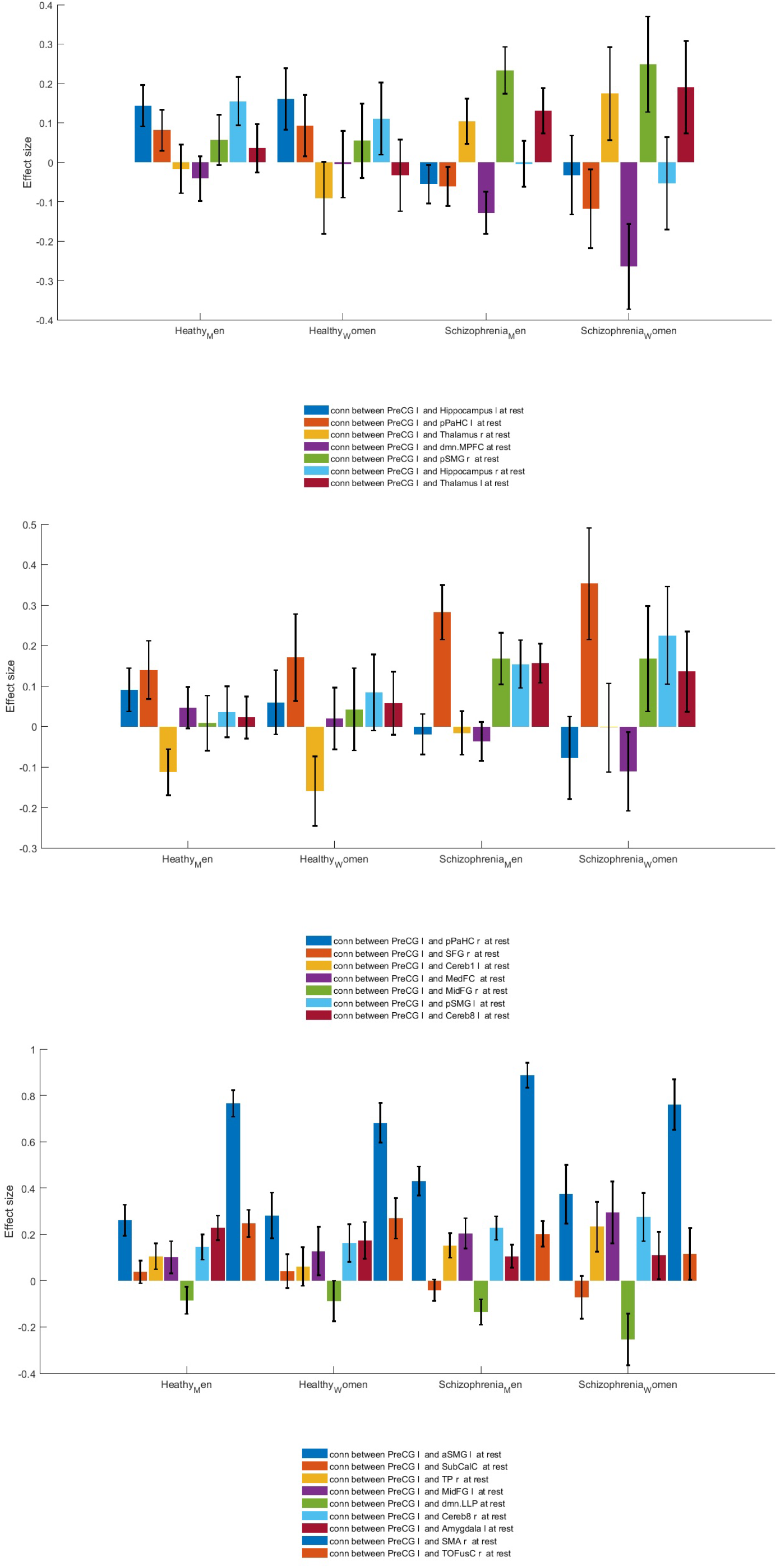
Functional connectivity difference of the … precentral gyrus between healthy controls vs. SCH patients (controls>Schizophrenia).

The functional connectivity (CONN) and Statistical Parametric Mapping (SPM) toolboxes [18, 5] used for spatially preprocessing (Realignment, coregistering, normalization) of structural and functional data of functional data is done using SPM toolbox. The functional volumes are coregistered with the region of interest and structural volumes. ROI-to-ROI correlational analysis were carried out by the CONN toolbox and SPM8. The preprocessing of the functional images considered of band-pass filtering of 0.008 - 0.09 Hz, motion correction, registration to structural images and spatial normalization to the MNI template. Then to reduce the physiological noise source, a Component Based Noise Correction Method (CompCor) has been used [2]. CompCor can be used for the reduction of noise in both blood oxygenation level dependent and perfusion-based functional magnetic resonance imaging data. False discovery rate correction is used to multiple hypothesis testing. Number of PCA components to be extracted for each ROI is set to one. It means that the time-series of interest is defined as the average BOLD activation within the ROI voxels, but it’s possible to define it as the principal eigenvariates of the time-series within the ROI voxels. Regions of interest and all the Brodmann areas defined from Talairach daemon assigned to all subjects. By segmentation of structural image for each subject, grey matter, white matter and cerebrospinal fluid (CSF) masks were generated.

Bivariate correlation is used as a functional connectivity measures between two areas. General linear model (GLM) [12] used for comparison of connectivity results between genders and between control and Schizophrenia patients.

### 4.1 ROI based analysis

We investigated the hypothesis of connectivity differece in schizophrenia and used ROI analysing for the PreCG region of the brain. Two-sample t-test analyses computed via SPM8 [6] to compare the connectivity results of patients vs. controls and male patient vs. women patient to compare the connectivity across two group. Connectivity values (Fisher-transformed correlation coefficients) between the seed and the identified ROI was extracted from the connectivity map. Different source of interest can be defined for analyzing.

### 4.2 First level voxel-based analysis

For a subject or condition it is possible to perform voxel-to-voxel analyzing that applies matrix of voxel-to-voxel connectivity values and there is no need for priori region of interest or seed analysis. In this method we can investigate whole brain connectivity. The voxel based analyzing can be based on connectivity pattern (Principal Component Analysis) between a voxel and the rest of the brain (*MVPA*). Another voxel based measeure is avialable in CONN toolboxes is *Indexes* that calculates the the average local connectivity between each voxel and its neighbors (Integrated Local Correlation) [4] or instead of average, the spatial asymmetry of the local connectivity can be used (Radial Correlation Contrast) [7]. Also, instead of local connectivity, global connectivity pattern between a voxel and the rest of the brain can be used (e.g. Radial Similarity Contrast) [9]. More details about measuring the Index can be found in [18].

## 5 Second level analysis

In the second level analysis step the between-subject contrast can be consider (e.g. to compared different groups like male vs. femals to see main effects in the connectivity within each group). In the ROI-to-ROI analyses, the first-level connectivity-measure matrix is used and the results can be thresholded at the desired p-value threshold. In this step by graph theoretical analyzing method provides the network measures like efficiency, centrality, and cost/degree to test the between-subject contrast.

## 6 Results

The connectivity contrast values between the PerCG right and the targets shows that the PreCG has abnormal communication with Thalamus, Hippocampus, Parahippocampal Gyrus (pPaHC), posterior division of Supramarginal Gyrus (pSMG) and medial prefrontal cortex (mPFC) (p-value=0.05). Figure 6 shows ROI-level results for PrecG left and right PreCG. Effects size for each region is shown by the dot size.

Functional connectivity difference of the left and right PreCG between healthy controls and SCH patients (controls>Schizophrenia) shows that the Connectivity of the left Hippocampus with both left and right PreCG decreased in SCH. More details about the the connectivity impairment of PreCG and the rest of the brain is demonstrated in figure 6. We also compaired SCH effects the PreCG in the mail and female patients (figure 6).

## 7 Conclusion

Here, we investigate abnormal connectivities of left and right precentral gyrus in male and female Schizophrenia patients. Region of interest based analysis carried out for the healthy control and Schizophrenia patient data. To do this resting state functional magnetic resonance imaging data of healthy control subjects and SCH patients from Center for Biomedical Research Excellence are used to examine the aberrant functional brain connectome in Schizophrenia which contains raw anatomical and functional MR data from 72 patients with Schizophrenia and 75 healthy controls, ranging in age from 18 to 65 years old. We can summarize our results as follows:

- The precentral gyrus has abnormal communication with Thalamus, Hippocampus, Parahippocampal Gyrus (pPaHC), posterior division of Supramarginal Gyrus (pSMG) and medial prefrontal cortex (mPFC) (p-value=0.05).
- The right precentral gyrus has unusual connectivities with Hippocampus l, pPaHC l Thalamus l, Thalamus r and pSMG r.
- The right precentral gyrus has unbalanced connectivities with Hippocampus l, pPaHC l, Thalamus r, MPFC, pSMG r, Hippocampus r and Thalamus l.
- The connectivity impairments of the PreCG in SCH are different for male and female patients.

This is information is expected to provide a better Understanding about altered functional functional Connectivity of Primary Motor Cortex in Schizophrenia.

**Table 3:**
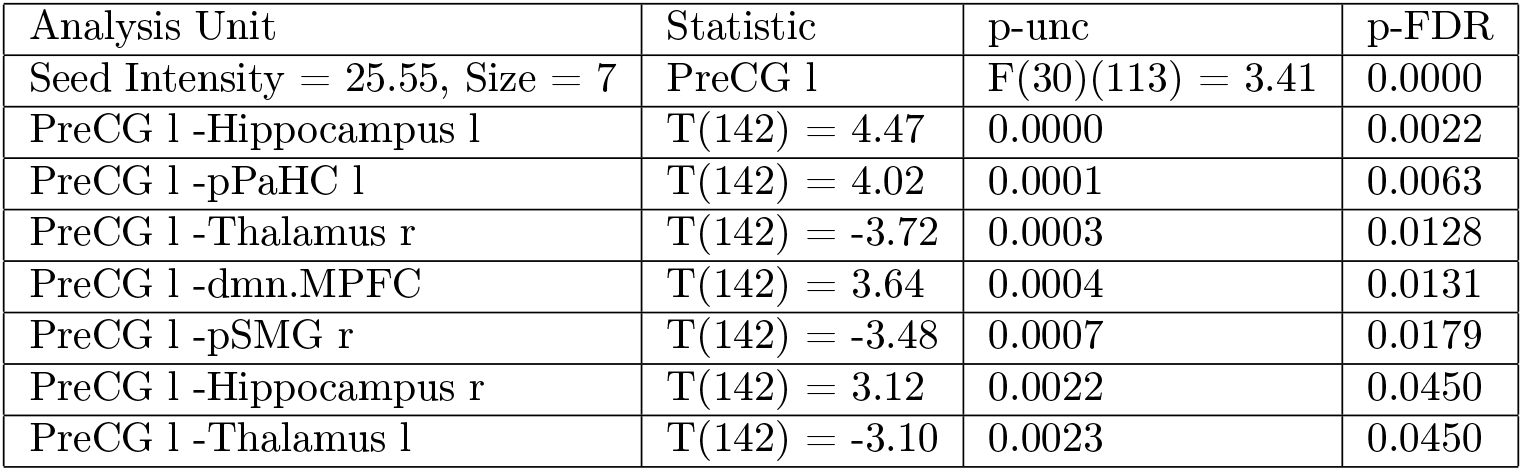
The connectivity contrast values between the PerCG left and the targets (p-value=0.05) (the complete list of the effect size is provided in the supplementary data)

